# The Unity/Diversity Framework of Executive Functions in Older Adults

**DOI:** 10.1101/2024.07.22.604231

**Authors:** Sheng-Ju Guo, Ping Wang, Lizhi Cao, Hui-Jie Li

## Abstract

Executive functions (EFs), encompassing inhibition, shifting, and updating as three fundamental subdomains, are typically characterized by a unity/diversity construct. However, given the dedifferentiation trend observed in aging, it remains controversial whether the construct of EFs in older adults becomes unidimensional or maintains unity/diversity. This study aims to explore and validate the construct of EFs in older adults. At the behavioral level, we conducted confirmatory factor analysis on data from 222 older adults who completed six tasks specifically targeting inhibition, shifting, and updating. One unidimensional model and six unity/diversity models of EFs were evaluated. Our results indicated that the EFs of older adults demonstrated greater congruence with the unity/diversity construct. At neural level, thirty older adults completed three thematically consistent fMRI tasks, targeting three subdomains of EFs respectively. Multivariate pattern analysis showed that rostromedia prefrontal cortex robustly showed similar neural representation across different tasks (unity). Meanwhile, the three EF domains were encoded by distinct global neural representation and the lateral prefrontal cortex play a crucial role in classification (diversity). These findings underscore the unity/diversity framework of EFs in older adults and offer important insights for designing interventions aimed at improving EFs in this population.

## Introduction

Executive functions (EFs) refer to high-level cognitive control processes that regulate thoughts and behaviors in a goal-oriented manner (Diamond, 2013; Miyake & Friedman, 2012), which are essential in our daily lives (Engel-Yeger & Rosenblum, 2021; Idowu & Szameitat, 2023; Smith et al., 2023). Over the past half-century, many researchers have employed various tests to explore the construct of EFs, yet no consensus has been reached (Baddeley & Hitch, 1974; Eslinger, 1996; Karr et al., 2018, 2022). The primary reasons stem from three aspects: the lack of a clear definition of EFs (Maldonado et al., 2020), the complexity of EFs measurement due to task impurity (Miyake & Friedman, 2012), and the variability in EFs performance across different age stages (Karr et al., 2022).

Given these considerations, Miyake et al. (2000) employed latent variable analysis at the behavioral level in young adults and found that EFs comprise three subcomponents (inhibition, shifting, and updating). Subsequent findings showed that the relationships among the three subcomponents can be classified into two behavioral models: the unidimensional/one-factor model and the multidimensional models (including a nested factor model, a three-factor model, three two-factor models that merged two EF subcomponents, and a bifactor model with a common EF factor and three specific factors) (Karr et al., 2018). The multidimensional models are described as “the unity/diversity framework”, which means that different EFs are correlated with each other (unity) but also show some separability (diversity) (Friedman et al., 2008; Miyake & Friedman, 2012).

The relationships among EF subcomponents may vary depending on the developmental stage of the individual. Specifically, EFs initially present as a unidimensional construct and progressively differentiate into a multidimensional construct throughout development (Karr et al., 2022). Previous studies have found that a unidimensional model is observed in early childhood samples (Masten et al., 2012; Wiebe et al., 2008, 2011; Willoughby et al., 2012), while three-factor models (Klauer et al., 2010; Miyake et al., 2000; Xu et al., 2013) and the nested factor model (Fleming et al., 2016; Friedman et al., 2008, 2009, 2011, 2016; Gustavson et al., 2022; Ito et al., 2015) are observed in adolescence and adulthood. However, the specific construct of EFs in older adults remains controversial. For instance, de Frias et al. (2009) found that EFs in older adults presented as a unidimensional model, potentially due to the increasing dedifferentiation of EFs with age, leading to enhanced correlations among EF subcomponents. Conversely, other studies supported two-factor or three-factor models of EFs in older adults (Karr et al., 2018; Ma et al., 2023), providing evidence for the unity/diversity construct of EFs. Previous behavioral studies on the EF construct of older adults have typically verified a particular conceptual model through confirmatory factor analysis (CFA). There is a lack of direct comparison among different models, making it difficult to determine which EF model is most suitable for older adults, leading to inconsistent findings regarding the construct of EFs in this population.

In addition to behavioral evidence, functional neuroimaging techniques can provide crucial insights into the construct of EFs at the neural level and further deepen our understanding of its underlying mechanisms. Multivariate pattern analysis (MVPA) is considered a promising method to explore the neural representations among different EF domains and may play a crucial role in validating the construct of EFs (Sandhaeger & Siegel, 2023). From a unity perspective, Vermeylen et al. (2020) conducted an impressive fMRI study that combined cognitive and affective domains to identify underlying shared neural patterns, finding universal representations within the medial frontal cortex. On the other hand, researchers are also committed to decoding the diverse neural representations in different executive control processes. They apply MVPA to investigate how the brain effectively distinguishes different conflict control processes in inhibitory control (Ritz & Shenhav, 2024), how it flexibly represents various task rules in shifting (Weber et al., 2023), and how it achieves selective representation and continuous updating in working memory (Oh et al., 2019). Recently, He et al. (2021) integrated the three subdomains of EFs and used MVPA to simultaneously reveal the unity and diversity of neural representation patterns in young adults. However, the evidence for the integration of EF representation patterns in older adults is still insufficient.

In the present study, we aimed to explore EF construct in older adults and validate whether the unity/diversity framework of EFs is preserved within this population. Specifically, we initially used tasks from three EF domains (inhibition, shifting, and updating) and employed CFA to compare seven EF models at the behavioral level. Figure 1 illustrates the seven behavioral EF models included in the present study. Subsequently, we conducted three gamified EF tasks and combined them with MVPA to verify the unity/diversity framework of EFs at the neural level. We first adopted searchlight analysis to identify brain regions exhibiting similar neural representations (unity) and then applied a three-way classifier to distinguish the global activation patterns of the three EF tasks (diversity).

**Figure 1.**
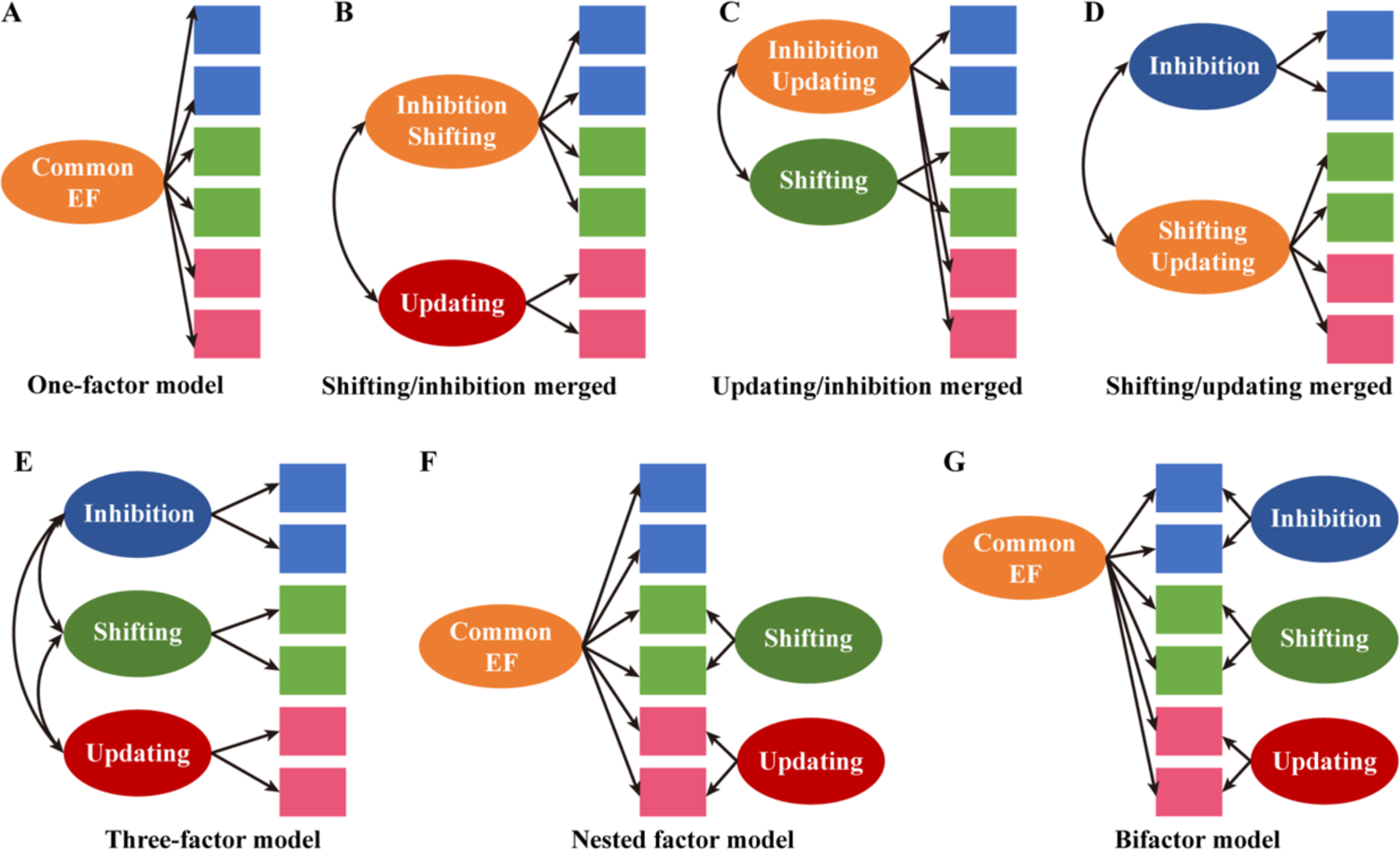
Overview of Factor Models Tested in Behavioral Analysis. (A) Unidimensional model. All the executive functions merged into one factor. (B) Shifting/inhibition merged model. The shifting factor and inhibition factor were merged into one factor and combined with the updating factor to construct the two-factor model. (C) Updating/inhibition merged model. The updating factor and inhibition factor were merged into one factor and combined with the shifting factor to construct the two-factor model. (D) Shifting/updating merged model. The shifting factor and updating factor were merged into one factor and combined with the inhibition factor to construct the two-factor model. (E) Three-factor model. Inhibition, updating, and shifting composed a three-factor model. (F) Nested factor model. It contains a common executive function bifactor, with shifting-specific and updating-specific factors respectively loaded on their selected indicators and no inhibition-specific factors. (G) Bifactor model. It contains a common executive function bifactor with specific factors for inhibition, shifting, and updating. Seven alternative measurement models were analyzed for comparison by using confirmatory factor analysis.

## Results

### Behavioral construct of EFs in older adults

We first assessed the construct validity of the unidimensional model using CFA. The results showed that the one-factor model fit had a poor fit (*χ^2^* (9) = 90.13, *p* < .001; *χ^2^*/*df* = 10.01; RMSEA = .20; CFI = .73; NFI = .72), indicating that the one-factor model did not fit our data. We then assessed the unity/diversity models. The fit indices revealed that the three-factor model had a good statistical fit to the data (*χ^2^* (6) = 5.02, *p* = .54; *χ^2^*/*df* = 0.84; RMSEA = 0.00; CFI = 1.00; NFI = 0.98), indicating that the hypothesized and real EFs construct were similar (Figure 2A). All the factor loadings and correlations between the latent variables (updating, shifting, and inhibition) in the three-factor model were significant (*p* < .01), ranging from moderate to high (0.40 to 0.88). The model fit indices of the nested factor model (*χ^2^* (5) = 7.31, *p* = .20; *χ^2^*/*df* = 1.46; RMSEA = 0.05; CFI = 0.99; NFI = 0.98) and bifactor model (*χ^2^* (3) = 5.02, *p* = .17; *χ^2^*/*df* = 1.67; RMSEA = 0.06; CFI = 0.99; NFI = 0.98) met the criteria. However, neither model converged and the estimated variances was negative, indicating that these two models did not fit the data. For the three two-factor models, the results showed that while the shifting/updating merged model (*χ^2^* (8) = 84.42, *p* = .00; *χ^2^*/*df* = 10.55; RMSEA = 0.21; CFI = 0.75; NFI = 0.74) and the updating/inhibition merged model (*χ^2^* (8) = 73.85, *p* = .00; *χ^2^*/*df* = 9.23; RMSEA = 0.19; CFI = 0.78; NFI = 0.77) did not meet the fit criteria, the shifting-inhibition merged model fit the data (*χ^2^* (8) = 7.39, *p* = .50; *χ^2^*/df = 0.92; RMSEA = .00; CFI = 1.00; NFI = 0.98). For the shifting/inhibition merged model (Figure 2B), all factor loadings and correlations between the latent variables were significant (*p* < .01). The correlation coefficient between two latent variables (updating, shifting-inhibition) was .50. Further analysis showed that the shifting/inhibition merged model had the equally good fit to the data as well as the three-factor model (*χ^2^_diff_* (2) = 2.38, *p* = .30). This suggested that both the three-factor model and the two-factor (shifting-inhibition, updating) model fit the EF constructs of older adults.

**Figure 2.**
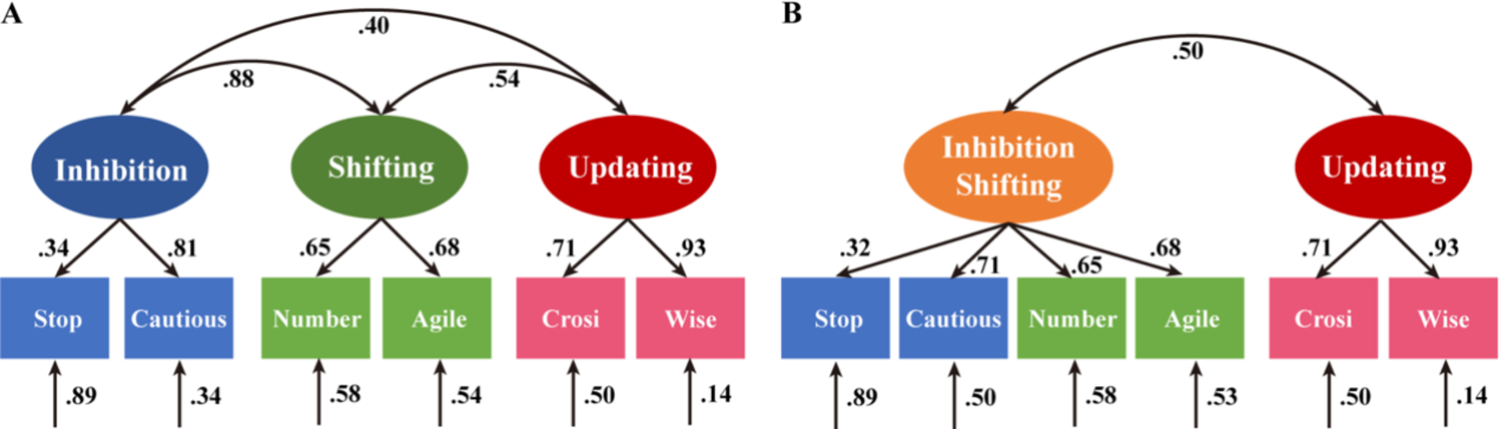
Parameterizations of Two Good-Fit Executive Function Models. (A) Three-factor model. The three factors (updating, shifting, and inhibition) are strongly correlated with each other (unity) but are separable (diversity) in that those correlations are far from 1.0. (B) Shifting/inhibition merged model. The merged factor (shifting-inhibition) and updating factor were moderately correlated, with correlations significantly greater than zero (unity) but also less than 1 (diversity). The numbers on the arrows are standardized factor loadings, those under the smaller arrows are residual variances, and those on curved double-headed arrows are interfactor correlations. The numbers in brackets are standard errors. All parameters were statistically significant (*p* < .05). Stop = stop-signal task; Cautious = Cautious Fisherman; Number = Number switch task; Agile = Agile Fisherman; Crosi = Crosi block-tapping task; Wise = Wise Fisherman.

In such correlated factor models, unity and diversity are reflected in the magnitudes of the correlations: factor correlations larger than zero suggest some unity, and correlations smaller than 1.0 suggest some diversity. To further validate the unity and diversity among different EF subdomains in older adults, we fixed the correlations between latent variables at either 0 or 1 (no unity or diversity, respectively) to assess whether there were significant changes in model fit. For the three-factor and two-factor (shifting-inhibition, updating) models, these comparisons indicate that the correlations between the latent variables are significantly greater than zero (three-factor model: *χ^2^_diff_* (3) = 125.30, *p* < .001; two-factor model: *χ^2^_diff_* (1) = 38.42, *p* < .001), and significantly less than 1.0 (three-factor model: *χ^2^_diff_* (1) = 86.97, *p* < .001; two-factor model: *χ^2^_diff_* (1) = 63.05, *p* < .001). Hence, each pair of latent variables was significantly correlated, but no pair was perfectly correlated, suggesting both unity and diversity among the different subdomains of EFs.

### Neural construct of EFs in older adults

#### Unity of neural patterns in EFs

The same analysis pipeline was replicated using searchlight analysis with different parameters (radius 3 mm and 6 mm). When the radius is 3 mm, brain regions showing similar neural representations were mainly located in the left rostromedial prefrontal cortex (RMPFC), right inferior parietal lobule, dorsal midcingulate cortex, and insula cortex (Figure 3A). Searchlight analysis with 6 mm radius further confirmed that the left RMPFC showed a greater ICC value (0.82 ± 0.02) (Figure 3B).

**Figure 3.**
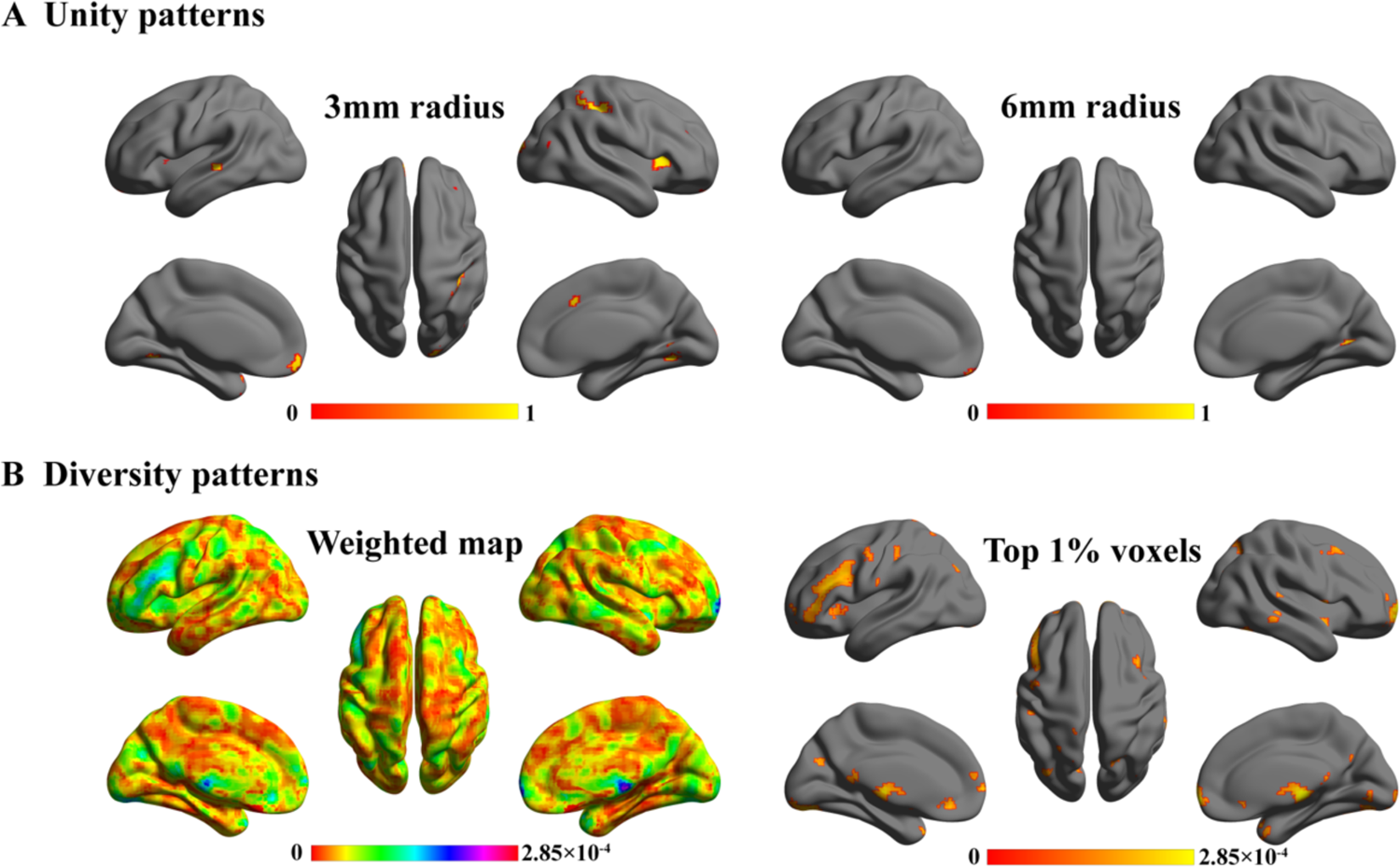
The Unity and Diversity of Neural Pattern Among Executive Function Tasks. (A) The results of searchlight analysis with radii of 3 mm and 6 mm are shown. Only voxels with intraclass correlation coefficient values greater than 0.80 are displayed. (B) The left panel shows the whole-brain weighted map. The right panel shows the top 1% of voxels with the highest weights (absolute value).

#### Diversity of neural patterns in EFs

Employing leave-one-out cross-validation, the three-way classifier demonstrated remarkable classification accuracy (92.22%) in distinguishing three EF subdomain tasks based on global neural representations, which was significantly greater than chance level performance (1,000 permutation tests, *p* < .001). Figure 3B displays the whole-brain weighted map, where the top 1% of high-weighted voxels are predominantly located in the right rostrolateral prefrontal cortex (RLPFC), premotor cortex (PMC), left inferior frontal gyrus (IFG), anterior cingulate cortex, and insula cortex, alongside segments of the parietal and temporal cortices. The prominence of these voxels underscores their pivotal role in the classification process, indicating their significant contribution to distinguishing among different EF tasks.

## Discussion

This study validated the construct of EFs in older adults at both behavioral and neural levels. At the behavioral level, the CFA supported both the three-factor model and the two-factor model (shifting-inhibition, updating) of EFs. These findings were further confirmed by comparing with no unity or diversity model, supporting the unity/diversity construct of EFs in older adults at the behavioral level. At the neural level, whole-brain MVPA was used to identify the representations of unity and diversity in EFs. Regarding the unity representation, the results revealed that the RMPFC robustly exhibited similar neural representations across different tasks. For the diversity representation, we found that distinct global neural representations encode inhibition, shifting, and updating, with extensive regions of the frontal and parietal lobes playing crucial roles in classification.

Previous behavioral studies have typically examined the construct of EFs in older adults by confirming a specific model through CFA (Adrover-Roig et al., 2012; de Frias et al., 2006; Vaughan & Giovanello, 2010; Wang et al., 2023a). Few have systematically compared all potential models (Karr et al., 2018, 2022) as the current study did, which has impeded the identification of the most suitable EFs construct for older adults. The current CFA results revealed that the construct of EFs was more congruent with the unity/diversity framework by simultaneously comparing the seven EF models proposed by previous studies (Karr et al., 2018). Specifically, the construct of EFs in older adults fit well with three-factor model and this diversity of the construct was further confirmed by comparing with perfectly correlated model. This finding is consistent with studies conducted on young adults (Fleming et al., 2016; Karr et al., 2018, 2022; Xu et al., 2013), suggesting that older adults reserved similar construct of EFs as young adults. In addition to the three-factor model, our results found that the two-factor model, in which shifting and inhibition were merged into one factor and updating as a separate factor, also had a good fit. This pattern potentially indicated a decrease in diversity between shifting and inhibition, reflecting a dedifferentiation trend in older adults (Koen et al., 2020; Koen & Rugg, 2019). Considering that our sample primarily consisted of young-old adults (Glisky et al., 2021), the good fit of the three-factor and shifting/inhibition merged models may indicate that older adults at this stage are in a transitional period of EF construct (Tang et al., 2023), during which young-old adults reserve the independence of various cognitive processes while also experiencing dedifferentiation due to aging (Koen et al., 2019).

Dedifferentiation preferentially occurs in similar cognitive processes (Ekstrom & Hill, 2023). According to Diamond (2013), shifting involves flexibly adaptation to new demands and rules, which inherently requires inhibition of previous tasks, rules, or perspectives. Previous studies have shown that among the three EF components, shifting is the first to decline (Diamond, 2013; Karr et al., 2022), causing older adults relying more on inhibition to complete shifting tasks. This reliance contributes to the dedifferentiation of the two components and the observed overlapping construct. Updating is typically associated with domain-specific processes such as rehearsal, coding, and storage (Frischkorn et al., 2022). These mental processes have little overlap with shifting and inhibition (Robinson & Steyvers, 2023), which may contribute to the maintenance of independence in the updating factor in older adults.

In addition to the behavioral analyses, the current study also examined the unity and diversity of neural representations of EFs, providing further support for the unity/diversity construct of EFs in older adults at the neural level. The searchlight analysis revealed that the RMPFC predominantly exhibited similar neural representations among three EFs components, indicating its primary role in the unity of EFs in older adults. This finding contrasts with a previous study in young adults, which reported the unity pattern of EFs located mainly in the lateral prefrontal cortex (LPFC) (He et al., 2021). This discrepancy can be explained by the Default-Executive Coupling Hypothesis of Aging (DECHA) (Spreng & Turner, 2019). According to DECHA, young adults mainly rely on the cognitive control and fluid abilities that are modulated by executive control network, specifically LPFC (Spreng & Turner, 2019; Stephenson et al., 2020; Tsumura et al., 2021). As people age, they tend to rely more on internal semantic knowledge to perform tasks, primarily due to the decline in cognitive control abilities. These changes lead to the semanticization of cognition and increase the coupling between the executive control network and default mode network (DMN) (Spreng & Turner, 2019). RMPFC is a crucial region within the DMN and plays a critical role in storing prior knowledge and experience (Jarovi et al., 2023; Rouse et al., 2024; Vaidya & Badre, 2022). The shift from LPFC in young adults to RMPFC in older adults, the shift in regions responsible for the unity of EFs may indicate this semanticization of cognition in older adults.

The three-way classifier demonstrated that the global neural representation could distinguish the three EF subdomains with high classification accuracy, indicating that EFs have distinct neural representation patterns throughout the brain and can efficiently represent and process different tasks. By quantifying the feature weights of each voxel, we observed that several regions with high weights, including PMC, RLPFC, and IFG, played important roles in domain-specific classification. RLPFC and PMC are typically characterized as domain-specific regions (Badre & Nee, 2018; Levy, 2024). The domain-specific characteristics of these two regions are further supported by the distinct neural representations observed in diverse EF domains in young adults (He et al., 2021), which is consistent with the current findings. Converging evidence from meta-analytic neuroimaging studies confirms the crucial domain-specific role of IFG (Narayanan et al., 2020; Turker et al., 2023). Badre and Nee (2018) proposed that the lateral frontal cortex exhibited hierarchical functional organization in EFs, with DLPFC as the apex, which is primarily repsonsible for higer-level abstract and general processing. In contrast, the posterior areas, including the IFG, are mainly responsible for lower-levels of concrete and specific processing in EFs. Overall, the current findings support the diversity of EFs in older adults, with both global and local neural representations contributing to distinguish different subdomains of EFs.

Limitations of the present study should be mentioned. First, the construct of EFs may be dynamic. The participants included in the current study were cross-sectional and had a relatively narrow age range. Future research is necessary to further investigate the age-related changes in the construct of EFs and the critical transition age in the aging process over a wider age range through a longitudinal design. Second, the present study included only healthy older adults. It is unclear whether the current findings can be generalized to older adults in clinical settings. Future studies are warranted to further confirm the unity/diversity construct of EFs in clinical individuals.

## Conclusions

The current study investigated the construct of EFs in older adults and found that it is more closely aligned with the unity/diversity framework by integrating multimodal data from behavior and brain imaging. These findings contribute to a more comprehensive understanding of the behavioral and neural mechanisms of EFs and also provide guidance for designing interventions aimed at improving EFs in aging population.

## Materials and Methods

### Participants

A total of 232 participants were recruited through online advertisements. None of them reported a clinical history or current diagnosis of neurodegenerative diseases (e.g., Alzheimer’s disease, Parkinson’s disease) or major psychiatric disorders (e.g., schizophrenia, depression, bipolar disorder). Other inclusion criteria included: age ≥ 60 years, years of education ≥ 6 years, right-handedness, normal or corrected-to-normal vision, and a Mini-Mental State Examination (MMSE) score of 24 or higher. After screening, 222 older adults (aged 66.76 ± 4.71 years, ranging 60 to 83 years; 163 females; Table 1) who met the inclusion criteria were included in the following CFA. The sample size was determined based on previous research regarding sample size requirements for structural equation models (Wolf et al., 2013).

**Table 1.** Demographic information for the two groups of older adults.

|  | Behavioral group | fMRI group |
| --- | --- | --- |
|  | (N = 222) | (N = 30) |
| Demographic variables | *M (SD)/%* | *M (SD)/%* |
| Age (year) | 66.76 (4.71) | 64.77 (3.94) |
| Female | 73.42 | 76.67 |
| Education (year) | 12.05 (2.85) | 13.13 (2.56) |
| MMSE (score) | 28.73 (1.17) | 28.73 (1.17) |

Note. MMSE = Mini-Mental State Examination

Among the aforementioned participants, 34 individuals without MRI contraindications (e.g., claustrophobia or metallic implants) were included in the fMRI validation study. During the fMRI scans, participants performed three EF tasks targeting inhibition, shifting, and updating, respectively. Three participants did not complete the shifting task, and one participant was excluded due to excessive head motion (≥ 3 mm). The final sample size was 30 (aged 64.77 ± 3.94 years, ranging from 60 to 74 years; 23 females).

The study was approved by the Ethics Committee of the Institute of Psychology, Chinese Academy of Sciences, and written informed consent was obtained from all participants before the study. Detailed demographic information of the participants is listed in Table 1.

## Measurements

### Executive function tasks in behavioral level validation

#### Inhibition

Inhibition was measured using the stop-signal task (Logan & Cowan, 1984) and Cautious Fisherman (Wang et al., 2023b). The commonly used metrics are stop-signal reaction time (SSRT) and accuracy (Friehs et al., 2020; Schmitt et al., 2016). Considering the speed-accuracy trade off, the task performance was measured using the composite Z score (sum of transformed Z of negative SSRT and accuracy) (Lyons et al., 2014).

#### Shifting

Shifing was evaluated using the number switch task (Schuch & Koch, 2003) and Agile Fisherman (Wang et al., 2023b). Shifting performance was assessed using the composite Z score (sum of the accuracy, negative repeat RT, and negative switch RT).

#### Updating

Updating was assessed using the Corsi block-tapping task (Kessels et al., 2000) and Wise Fisherman. The dependent variable is the Z score of total number of correctly clicked squares (Wang et al., 2023b).

### Executive function tasks in neural level validation

Considering the task-impurity caused by specific task context (He et al., 2021; Miyake & Friedman, 2012), FISHERMAN was used as the fMRI task, which contains three EF gamification tests with consistent thematic elements (Wang et al., 2023b; Wang & Li, 2024). In these gamified tests, a fishing net appeared in the center of an ocean-themed environment, with a fish and a shrimp randomly appeared in eight white bubbles surrounding the fishing net. Participants caught marine life within bubbles by pressing buttons on a response pad according to specific rules. All participants completed practice sessions using a simulated MRI scanner to familiarize themselves with the response pad and the task. When the accuracy reached 70%, a formal scan was conducted.

#### Cautious Fisherman

The task targets inhibition and employs a mixed design (Figure 4A). Two conditions (stop trials and go trials) are included in the task. In go trials, participants were instructed to ‘catch’ the marine creatures that matched the color of the fishing net displayed in the center. In stop trials, a black bubble appeared after the marine creatures were displayed, and participants were asked to stop catching any marine creatures. The initial presentation time of black bubble is 250 ms (stop-signal delay, SSD) after the appearance of marine creatures. If participants successfully inhibited on a stop trial, the SSD is increased by 50 milliseconds to make subsequent stop trials more difficult; if the participant failed to inhibit, the SSD is decreased by 50 milliseconds to make inhibition easier. The intertrial intervals varies from 500 ms to 1500 ms (mean = 1000 ms). The fMRI task includes two runs, each lasting 8 min 10 s. Each run contains 12 blocks with 10 s interblock intervals for a total of 120 trials. Each block lasts 30 s and contained 10 trials. In each trial, marine creature is displayed for 1000 ms, and participants have 2000 ms to respond. The ratio of stop trial to go trials is 1:3.

**Figure 4.**
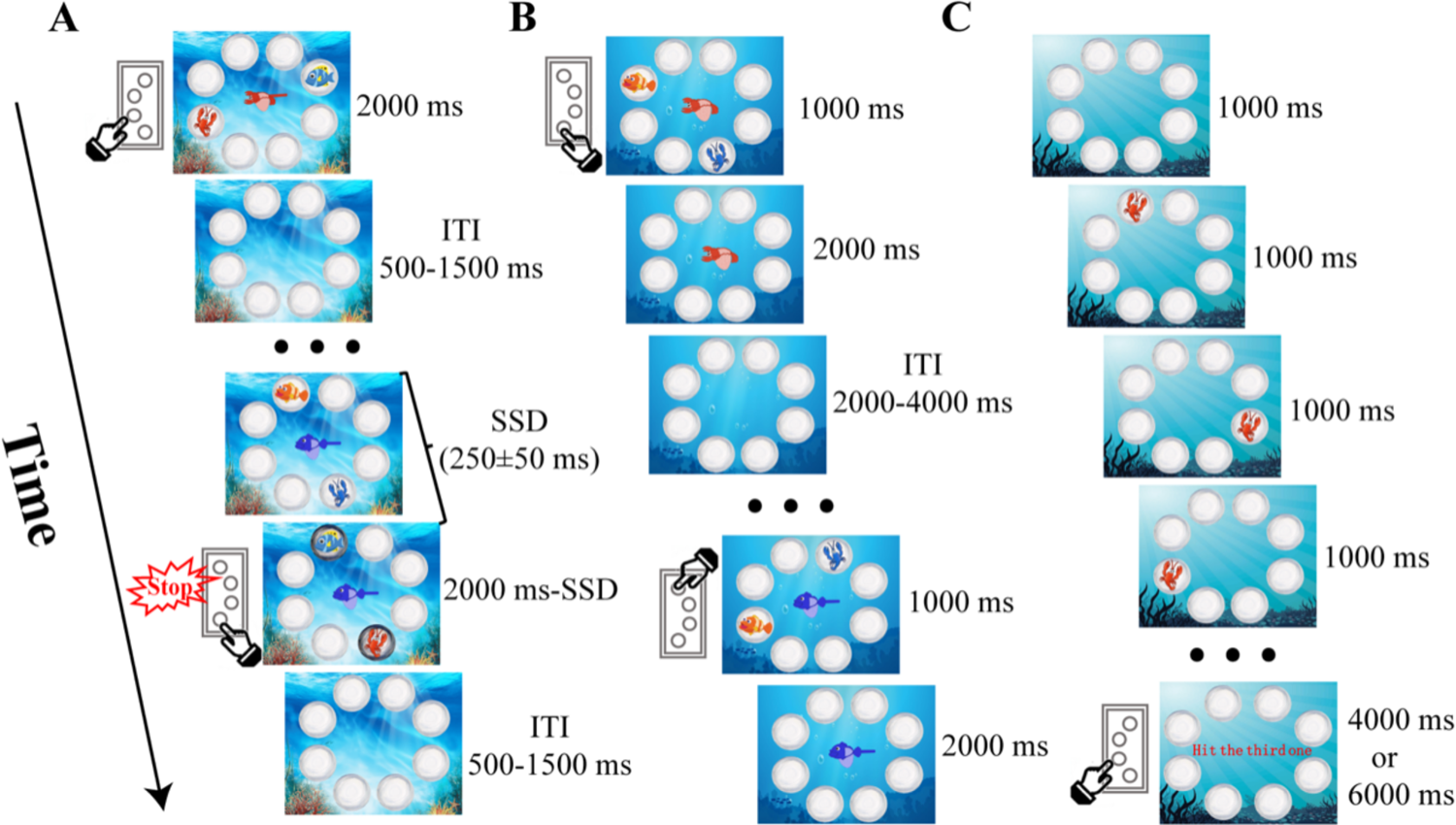
Flow Chart of the Functional Magnetic Resonance Imaging (fMRI) Paradigm of Three Executive Function Tasks. For the Cautious Fisherman (A), participants are required to catch marine creatures of the same color unless a black bubble appears, in which case they should withhold catching any creatures. For the Agile Fisherman (B), if the middle net has a handle, participants need to catch according to the color of the net; otherwise, they need to catch the creature with the same shape. For the Wise Fisherman (C), participants need to remember the order of creatures displayed and catch the creature according to the central instruction. There are 8 buttons on the response pad, corresponding to the bubbles in each direction. SSD = stop-signal delay; ITI = intertrial interval.

#### Agile Fisherman

The task targets shifting and employs an event-related design (Figure 4B) (Wang et al., 2024). The task consists of two conditions (switch trials and repeat trials). Each trial begins with eight blank bubbles as a baseline for 3000 ms, followed by the display of a fishing net and marine creatures for 1000 ms. Participants should respond within 2000 ms depending on the type of fishing net. They were instructed to catch marine creatures according to the color of the net (red or blue) when the net had a handle, or according to the shape of the net (fish or shrimp) when the net had no handle. The intertrial intervals varies from 2000 ms to 4000 ms (mean = 3000 ms). A trial (after the first one) is defined as a repeat trial if the type of the net is the same as the previous one, otherwise it is a switch trial. The fMRI task includes two runs of 8 min 10 s each, and each run consists of 80 trials. The two conditions are presented in a pseudorandom order with a 1:1 ratio.

#### Wise Fisherman

The task targets updating and uses a block design (Figure 4C). Two types of blocks, updating block and baseline block, were included in the task. The updating block lasted 20 s and consisted of two trials, 3 span and 5 span. In each trial, marine creatures are displayed one by one in random positions for 1000 ms each. Then, a response window appeared, during which participants had to recall and catch previously displayed marine creatures based on the central cue. The response window lasts 4000 ms for 3 span or 6000 ms for 5 span, with a 1000 ms interval between two trials. The baseline block lasts 10 s and participants are asked to stare at the screen with eight bubbles. This task contains two runs, each lasting 5 min 10 s. Each run consists of 10 updating blocks and 10 baseline blocks.

### Functional imaging data acquisition and preprocessing

The experiment was conducted on a 3T MRI scanner (GE Discovery MR750) at the Magnetic Resonance Imaging Research Center of the Institute of Psychology, Chinese Academy of Sciences. Foam pads were used to support the head and neck to minimize head motion and reduce cumulative head drift during scanning. Functional T2*-weighted images were acquired using gradient-echo echo-planar imaging (EPI) sequences in a top-down sequential order during the task with to the following parameters: repetition time (TR) = 2000 ms, echo time (TE) = 30 ms, flip angle (FA) = 90°, voxel size = 3.5 × 3.5 × 3.5 mm^3^, slice thickness = 3.5 mm, slice number = 37 slices, matrix = 64 × 64, and field of view (FOV) = 224 × 224 mm^2^.

The fMRI data were preprocessed using SPM12 software (https://www.fil.ion.ucl.ac.uk/spm/). Preprocessing steps included head motion correction, slice timing correction, normalization, and spatial smoothing with a Gaussian kernel of 7 mm full width at half maximum (Figure 5A). Participants with large head motions (≥ 3 mm) were excluded from further analyses.

**Figure 5.**
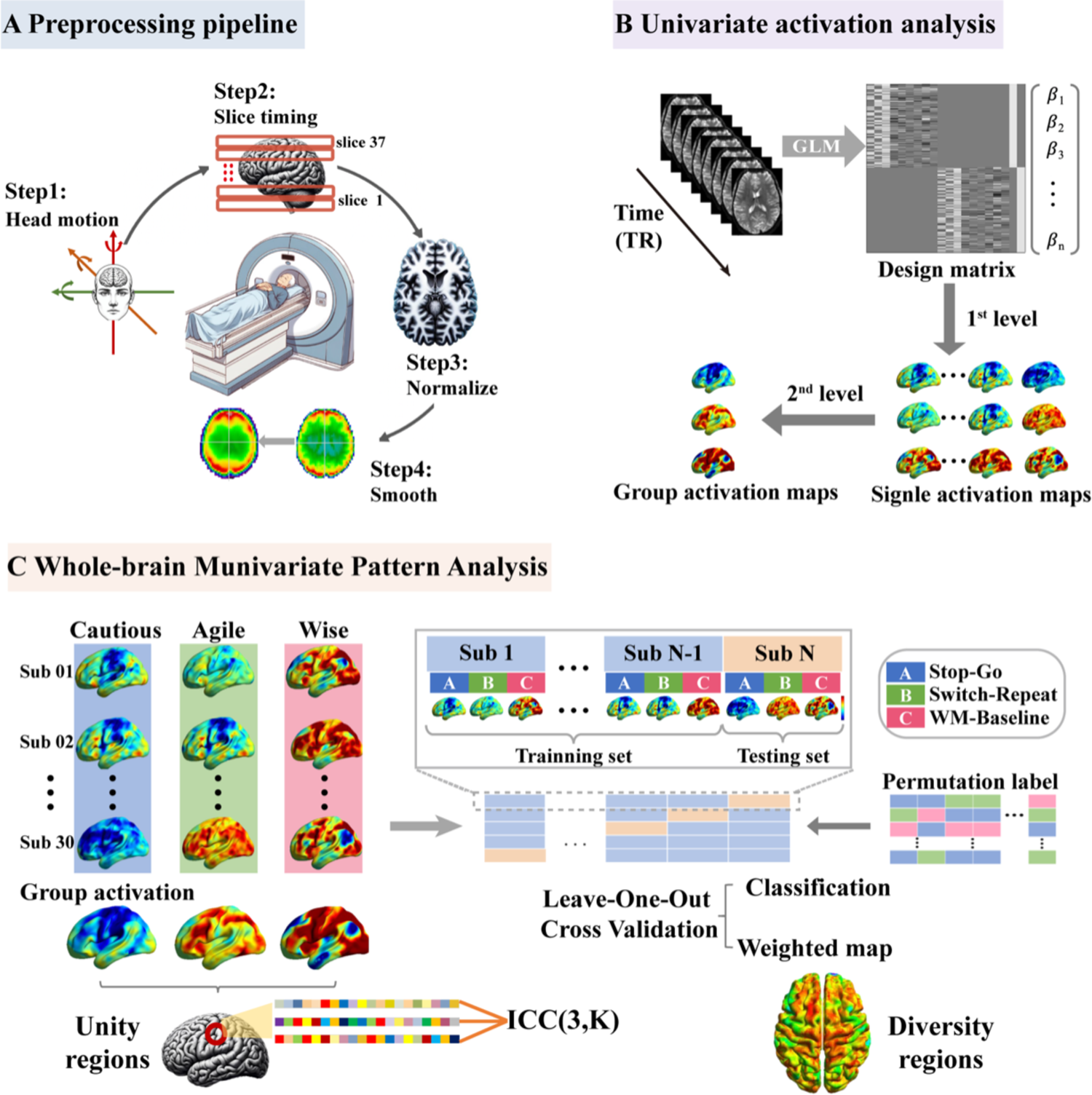
Schematic Overview of the fMRI Analysis. (A) Preprocessing pipeline. (B) Univariate activation maps for each task at the individual level and group level were obtained using first-level and second-level analysis in SPM12. (C) Whole-brain multivariate pattern analysis. Three group activation maps were used for searchlight analysis to identify unity regions based on the intraclass correlation coefficient. The β map from the three first-level contrast images for each participant was extracted as the model feature. The three-way classifier based on leave-one-out cross-validation computed the classification accuracy for discriminating the three EF tasks and further identified diverse regions based on a weighted map. A permutation test was performed to determine whether the accuracy was significantly greater than the values expected by chance. Cautious = Cautious Fisherman; Agile = Agile Fisherman; Wise = Wise Fisherman.

## Data analysis

### Confirmatory factor analysis

The construct of EFs in normal older adults was tested through CFA based on participants’ performance on each behavioral task. The subsequent comparative analyses used seven models, including a unidimensional model (Figure 1A), three two-factor models that merged two of the three factors (e.g., inhibition-updating; updating-shifting; inhibition-shifting) (Figure 1B, 1C and 1D), a three-factor model (e.g., inhibition, updating, and shifting) (Figure 1E), a nested factor model (e.g., common EF, shifting-specific and updating-specific factors) (Figure 1F), and a bifactor model (e.g., common EF, inhibition-specific, shifting-specific, and updating-specific factors) (Figure 1G). Statistical analysis was performed using the package lavaan in R (Rosseel, 2012).

The model validation was examined through the following two steps. The first step was to ensure proper model convergence, meaning that the model converged without any errors that indicate an unacceptable solution or unreliable estimates (e.g., correlations with absolute value greater than 1.0, negative residual variances, nonpositive definite latent variable covariance matrices). If the model is converged, the model fit is evaluated by following indices: chi-square (*χ*^2^), normed chi-square (*χ*^2^/df), comparative fit index (CFI), normed fit index (NFI) as well as root mean square error of approximation (RMSEA). The criteria for good model fit are nonsignificant *χ*^2^, *χ*^2^/df < 2, CFI > .90, NFI > .90, and RMSEA < .06 (Byrne et al., 1989; Hu & Bentler, 1999).

The second step was model comparisons. Models meeting these criteria are included in chi-square difference (*χ*^2^*_diff_*) tests to select the best-fitting model (Friedman et al., 2011).

### fMRI univariate activation analysis

Univariate activation analysis was performed using SPM12 (Figure 5B). To calculate the blood oxygen level-dependent (BOLD) signal under different conditions of the three tasks, a general linear model (GLM) was used for the first-level analysis. The regressors for the Cautious Fishermen task included go condition, stop condition, and baseline. The regressors for the Agile Fishermen task included the switch condition, repetition condition, and baseline. The regressors for the Wise Fishermen task included 3 span condition, 5 span condition, and baseline. Each model included six raw head motion parameters as covariates to eliminate motion. All regressors were convolved with the canonical hemodynamic response function. A high-pass filter of 1/128 Hz was implemented to remove low-frequency drift from the time series. Contrasts were set for each model, including the stop condition versus the go condition for the Cautious Fishermen task, the switch condition versus the repeat condition for the Agile Fishermen task, and the updating condition versus the baseline for the Wise Fishermen task. These first-level contrasts were submitted to second-level analysis by one-sample t-test to identify global brain activation when performing the task. The results of the first- and second-level analyses were then used for subsequent MVPA.

### Whole-brain multivariate pattern analysis

To validate the unity/diversity framework of EFs at the neural level, we conducted searchlight MVPA and whole-brain classification analyses on different EF tasks (Figure 5C). Surface mapping was conducted using BrainNet Viewer (Xia et al., 2013) to visualize the results on the cortical surface.

#### Searchlight MVPA

Searchlight MVPA was performed using the CoSMoMVPA toolbox (Oosterhof et al., 2016; http://cosmomvpa.org/) to identify regions with similar neural representations in the three EF subdomains across the whole brain. First, we defined a small sphere with a radius of 3 mm (Axelrod et al., 2017). To ensure the stability and reliability of the results, a larger sphere with a radius of 6 mm was additionally used for validation (Etzel et al., 2013; Kriegeskorte et al., 2006; Oosterhof et al., 2016). For each sphere, we extracted three vectors of parametric values from the three second-level contrast images and calculated the intraclass correlation coefficient (ICC), which serves as an estimate of local neural representation similarity. We focused on ICC values greater than 0.80, a threshold indicating high reliability and consistency (Landis & Koch, 1977).

#### Whole-brain classification analysis

Whole-brain classification analysis was used to examine whether there were different global neural representations among the three EF tasks. This analysis consisted of two steps. First, the estimated first-level brain maps of the three tasks were normalized and used as input features for subsequent analyses. Then, we applied a global-level classifier to decode the three task contrasts using the Nilearn package (Abraham et al., 2014; https://nilearn.github.io/). A three-way classifier was trained using linear support vector machines with default parameters (penalty = l2). Leave-one-out cross-validation was used to evaluate the classification performance of neural representation patterns associated with different tasks, measured by the average classification accuracy. To improve the robustness of the results, each cross-validation was repeated ten times, generating reliable measures of classification accuracy (Valente et al., 2021). We also implemented a permutation test in which task labels were randomized 1,000 times and replicated the MVPA pipeline to ensure that the accuracy was significantly higher than the chance level (He et al., 2021). Significance was statistically inferred from the null distribution of accuracy, calculated as the proportion of permutations that produced accuracy greater than the actual value. The above classification analysis generated three classifiers with different feature weights, representing the neural activity patterns associated with specific tasks. The higher the absolute value of the weights, the greater the contribution of the corresponding feature to the classification. Therefore, we calculated the average absolute weight of each voxel to identify different representation regions.

## Impact Statement

The executive functions of older adults exhibited a unity and diversity construct, with the former mediated by the rostromedia prefrontal cortex and the latter by the lateral prefrontal cortex.

## Acknowledgments

We thank all the participants who participated in this study.

## Funding

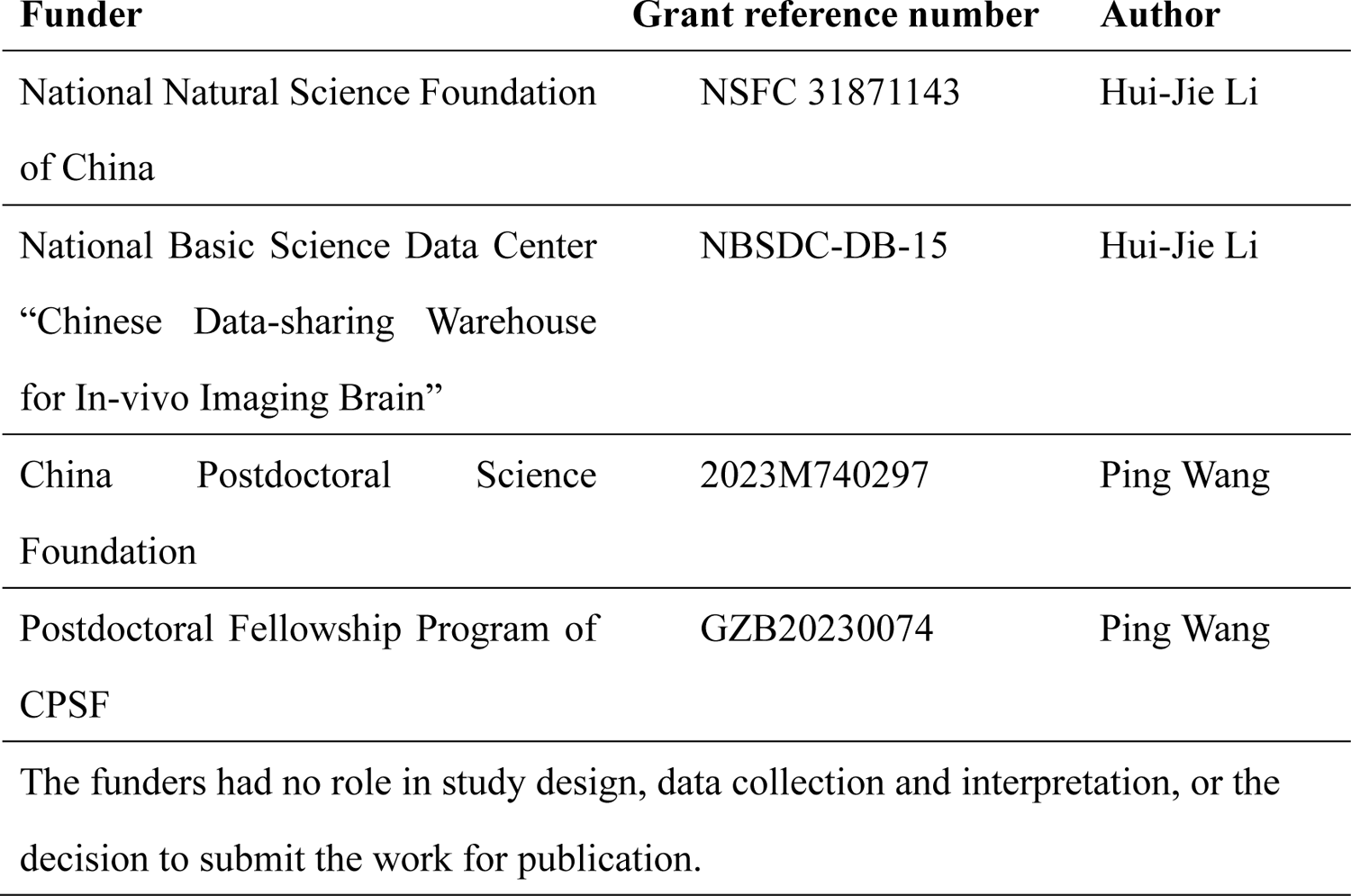

## Author contributions

Sheng-Ju Guo, Ping Wang, Data curation, Formal analysis, Validation, Investigation, Visualization, Methodology, Writing - original draft; Hui-Jie Li, Conceptualization, Supervision, Writing – review and editing

## Ethics

Human subjects: The experiments were approved by the Institutional Review Board of the Institute of Psychology, Chinese Academy of Science (Approval #H21015). Informed consent was obtained from all subjects.

